# Incremental AMR acquisition driving successive genotype replacements and the rise of extensively drug resistant (XDR) *Shigella sonnei* in Australia over 20 years

**DOI:** 10.64898/2026.06.17.728071

**Authors:** Jake A Lacey, Sally Dougall, Mathilda Wilmot, Himal Shrestha, Karolina Mercoulia, Paula Roydhouse, Jessica Barnden, Susan A. Ballard, Mihaela Ivan, Christian McGrath, Danielle J Ingle, Benjamin P Howden, Norelle L Sherry

## Abstract

**Background:** In Australia, the burden of shigellosis is predominantly in returning travellers or in men who have sex with men (MSM). Here, we combine genomic data with comprehensive epidemiological data on sexual exposure and international travel to explore population dynamics of *Shigella sonnei* and the expansion of multi-drug resistant (MDR) and extensively drug-resistant (XDR) sub-lineages.

**Methods:** A population-level study of all cultured *Shigella sonnei* isolates in the state of Victoria, Australia, was undertaken between January 2002 and December 2024. Antimicrobial susceptibility testing, whole-genome sequencing, and bioinformatic analyses of 1,305 *Shigella sonnei* isolates were performed at the Microbiological Diagnostic Unit Public Health Laboratory. Enhanced metadata on source attribution including travel and sexual exposure were collected through surveillance forms or by interviews.

**Results:** This study highlights significant shifts in *Shigella sonnei* cases in Victoria from sensitive strains to MDR and then XDR, particularly in the MSM-associated groups but also associated with a large point source outbreak. We describe an historical pattern of shifting genotype prevalence, and replacement to more varied and higher proportions of antimicrobial resistance over the last decade, resulting in the establishment of two distinct but highly concerning XDR sub-lineages within Victoria.

**Conclusions:** Our genomic-epidemiological analyses highlight that drug-resistant *Shigella sonnei* remains an ongoing public health threat, and the importance of ongoing surveillance. We determined local evolutionary trajectories and identified expanding sub-lineages that informed shifts in clinical management and antimicrobial recommendations over time, including the use of azithromycin and carbapenems. Placing these local dynamics within the broader global epidemiology, we link how regional evolution interconnects with international dissemination, proving valuable context for guiding local, national and global strategies for prevent outbreaks and antimicrobial resistance.

## Introduction

*Shigella <u>sonnei</u>* is one of four species of *Shigella,* a highly infectious enteric bacteria that cause shigellosis, a form of bacillary dysentery, and is transmitted primarily via faecal-oral contact ^1^. Globally, *Shigella* is consistently a top contributor to the gastrointestinal disease burden due to its high infection rates and severe health outcomes, especially in children from low- and middle-income countries^2^. In high-income countries (HICs) including Australia, point source outbreaks of shigellosis driven by *S. sonnei* and *S. flexneri* have been commonly linked to food sources or identified in returned travellers ^3–6^, while *S. boydii* and *S. dysenteriae* are comparably rare. In the past decade, there has been a significant change in the epidemiology of *S. sonnei,* with cases of endemic shigellosis in men in HICs now considered associated with sexual transmission among men who have sex with men (MSM)^7–11^.

Antimicrobial resistance has emerged as an issue in *Shigella* over the past decade, with increasing rates of multidrug resistance (MDR) and, more recently, extensively drug-resistant (XDR) *Shigella* being reported globally, with a notable rise in the past five years ^7,8,12–15^. Definitions of MDR and XDR vary between studies and surveillance systems. Generally, MDR refers to resistance to three or more antimicrobial classes^16^. The U.S. Centers for Disease Control and Prevention (CDC) define XDR *Shigella* as resistant to all commonly recommended antibiotics, including azithromycin, ciprofloxacin, ceftriaxone, trimethoprim-sulfamethoxazole and ampicillin.

Outbreaks of MDR and XDR *S. sonnei* are being increasingly reported in high-income regions predominantly affecting MSM communities^10,12,17–19^. Antimicrobial resistance (AMR) in *Shigella* is frequently plasmid-mediated ^13,20^, enabling rapid dissemination of resistance genes between strains and across species ^8,13,17^. This has facilitated the emergence of novel resistant lineages, including those carrying third-generation cephalosporin (3GC) resistance determinants^12,13^ that have spreads globally in both *S. sonnei* and *S. flexneri*^21^ species. This has resulted in subsequent clonal expansions of increasingly resistance lineages over time. Due to the clonal nature of *S. sonnei,* these features are only identifiable by genomic epidemiological surveillance to track transmission within and between countries.

These evolving AMR patterns pose increasing challenges for both clinical management and public health due to limited effective treatment options. Whilst treatment recommendations previously recommended ciprofloxacin as first-line therapy, increasing resistance in some countries has led to alternative agents such as azithromycin and ceftriaxone now being recommended depending on local susceptibility patterns, including Australia^22^.

Here we present a retrospective, observational study of 1,305 cases of *S. sonnei* between January 2002 and May 2024. We aim to understand and comprehensively characterise the shifting patterns of antimicrobial resistance and population dynamics of *S. sonnei* in Victoria, Australia using integrated genomic and epidemiologic data. The longevity and depth of this dataset provide unique and robust insights to enable us to investigate the longitudinal trends and evolutionary shifts within the population which shorter term studies cannot capture.

## Methods

### Study Setting

In Australia, shigellosis is a notifiable disease under public health legislation, and primary diagnostic laboratories are required to forward all *Shigella* isolates to the Microbiological Diagnostic Unit Public Health Laboratory for further characterisation, including phenotypic susceptibility testing and routine WGS. The Microbiological Diagnostic Unit Public Health Laboratory (MDU-PHL) is the bacteriology reference laboratory for the state of Victoria (population ∼6.97 million people in 2024). *Shigella* spp. identification at MDU-PHL from 2002-2024 was performed using a combination of serological and standard biochemical tests. In house biochemical tests based on the Ewing and Edwards typing scheme (1986) were used to confirm the isolate as *Shigella* spp. and serotyping performed by slide agglutination tests using commercial antisera to LPS O-antigen. Antisera were sourced from Denka Seiken co., LTD (Tokyo, Japan) and MAST Group LTD (UK). Polyvalent antisera confirmed the *Shigella* subgroup (A,B,C,D) and type specific monovalent antisera confirmed the serotype^23^.

### Whole-genome sequencing and Quality Control

DNA extractions were prepared using Illumina Nextera XT DNA library preparation and whole genome sequencing of study isolates were sequenced on Illumina platforms (Nextseq 500/550). Quality control including reads assessment, species identification, genome assemblies, annotation, and assessment of genome metrics was conducted using the bohra microbial genomics pipeline (https://github.com/MDU-PHL/bohra) using seqkit v2.1.0, KMC v3.2.1, Kraken v2.1.2, shovill v1.1.0, SPAdes v3.13.2, prokka v1.14.6. All read data for this project has been deposited in Bioproject PRJNA319594, and details of included isolates are provided in Supplementary Table 1.

### Genotyping

*In silico* species confirmation and serotyping was performed using ShigaPASS^24^ with the draft genome assemblies as input. *In silico* multilocus sequence typing (MLST) using the *E. coli* achtman_4 scheme (https://pubmlst.org/bigsdb?db=pubmlst_escherichia_isolates) was conducted to assign sequences a sequence type (ST) using mlst v2.23.0 (https://github.com/tseemann/mlst). Genotyping in previously characterised *S. sonnei* lineages was undertaken with SonneiTyping implemented in mykrobe v0.13.029 using the short-read data ^25,26^.

### Variant Calling and Phylogenetic inference

Identification of core genome single nucleotide polymorphisms (SNPs) against a reference genome AUSMDU00008333 (accession: SAMEA5210148) were performed using snippy v4.4.5 (https://github.com/tseemann/snippy, applying minfrac 10 and mincov 0.9). The SNP variants identified by snippy were used for phylogenetic reconstruction using maximum likelihood (ML) in IQtree2 v2.1.4^27^ with constant sites corrected, 1000 bootstraps, and a generalised time-reversable model of evolution GTR+F+G4. Treetime was utilised for ancestral state reconstruction and inference of timed tree phylogenies applying the -- covariation --stochastic-resolve --coalescent skyline –confidence parameters ^28^. Phylogenies and skyline plots were processed and visualised in R v4.3.1 using the ggtree v3.8.2, treeio v1.24.3, ggplot v3.5.0, dplyr v2.4.0, readr v2.4.1 and ape v5.7-1 packages.

### Timeline analysis and estimation of population shifts

To investigate the temporal patterns/trends of *S. sonnei* genotypes across the population, and MSM and travel-associated sub-populations, we performed a cumulative incidence analysis. We analysed monthly trends of *S. sonnei* genotypes and AMR profiles using R v4.3.1. The temporal analysis was restricted to *S. sonnei* isolates from 1 January 2015 to 11 November 2024 where month and year data were available for 1,261 isolates received from 2015. Data recorded before 2015 did not record MSM status as a risk factor. A Year Month column was generated by flooring dates to the start of each month. The ten most frequent genotypes were identified using count() and retained for visualization. All other genotypes (n = 24) were reclassified into a single group (“Other”) using R v4.3.1. Cumulative incidence curves were plotted to demonstrate the trends.

We performed a joinpoint regression analysis to identify temporal shifts in the incidence of *S. sonnei* genotypes in each risk group and the entire dataset. We grouped isolates into their respective genotypes and applied a joinpoint regression using the segmented v1.6-4 function to detected statistically significant temporal breakpoints, in the monthly incidence of each genotype group. Poisson generalised linear models (GLMS) were used as the base model for each group. The number of breakpoints was iteratively tested and the optimal model was selected based on the lower Akaike information criterion (ASIC) breakpoints were extracted and covered to calendar date using the earliest month as the time origin. Observed counts and model trends were plotted onto a line graph in R.

Data wrangling and visualization were performed with the tidyverse v2.0.0, lubridate v1.9.3, ggplot2 v3.5.0, and RColorBrewer v1.1-3 packages.

### AMR detection and definition of MDR and XDR

Antimicrobial resistance genes were detected from genome assemblies using abriTAMR v1.0.14 ^29^, and single-nucleotide polymorphism (SNP) based mutations were included using species flag --*Escherichia*. SNPs within the quinolone resistance determining regions (QRDR) for *S. sonnei* were additionally confirmed using sonneityper^25^. Determination of MDR and XDR isolates were determined based on a custom script (https://github.com/JA-Lacey/R_helper_scripts/blob/main/R_Antimicrobial_resistance_interpretation/Shigella_Define_AMR_status_to_CDC_classificaiton.Rmd) that parses the outputs of abriTAMR according to the CDC definitions. Following the CDC definitions^16^, MDR was defined as *in silico* detection of AMR to at least three classes of antibiotics, and XDR was defined as *in silico* AMR to all commonly recommended empiric and alternative antibiotics, ciprofloxacin (AMR determinant: QRDR 3-point or 1-point + *qnrS1/S13*), ceftriaxone (AMR determinant: *blaCTX-M*), azithromycin (AMR determinant: *mph(A), ermB*), trimethoprim-sulfamethoxazole (AMR determinant: *sul/dfrA*) and ampicillin (AMR determinant: *blaCTX-M*). For further discrimination we additionally split MDR status into two categories: MDR, and MDR with extended beta-lactam resistance.

Antimicrobial susceptibility testing (AST) and laboratory confirmation of minimum inhibitory concentrations (MICs) for *S. sonnei* isolates was conducted in our laboratory using different techniques over the study period. Agar dilution was conducted from (2005–2021) and broth microdilution (BMD; 2022–present). Agar dilution testing followed Clinical and Laboratory Standards Institute (CLSI) guidelines (CLSI, 2017) and included the following antimicrobials over the testing period: ampicillin, streptomycin, tetracycline, chloramphenicol, sulphathiozole, trimethoprim, kanamycin, nalidixic acid, spectinomycin, gentamicin, ciprofloxacin, cefotaxime, azithromycin, meropenem, and trimethoprim/sulfamethoxazole. Since 2022 BMD testing has been conducted according to EUCAST methods and interpreted according to EUCAST clinical breakpoints 2026v16 using the Sensititre NARMS Gram Negative CMV5AGNF antimicrobial susceptibility plate (ThermoFisher). Antimicrobials tested included cefoxitin, azithromycin, chloramphenicol, tetracycline, ceftriaxone, ampicillin/clavulanic acid, ciprofloxacin, gentamicin, nalidixic acid, meropenem, sulfisoxazole, trimethoprim/sulfamethoxazole, ampicillin, and colistin.

### International context isolates

To provide phylogeographic context for the Australian *S. sonnei* isolates, we uploaded all read sets from this study to the NCBI Sequence Read Archive (SRA) and integrated those datasets into the Pathogen Detection Portal (PDP). With the PDP framework genomes were subject to the NCBI SNP pipeline to define clusters (https://ftp.ncbi.nlm.nih.gov/pathogen/Methods.txt). Briefly the pipeline involves masking repeat regions, removing poor-quality assemblies, and partitioning isolates within each organism group using pairwise k-mer distances. Clusters are formed by single-linkage clustering with a k-mer distance threshold of 0.1 that are broad enough to retain closely related pairs, though isolates within a partition may still differ by thousands of SNPs. After refinement based on SNP comparisons to define target partitions, each partition is processed by selecting an internal reference to compute SNPs, generate pairwise SNP matrices, and produce maximum compatibility trees and associated outputs for public release. We searched all isolates to determine their corresponding cluster designations. For each identified cluster, we downloaded additional publicly available raw sequencing reads where key metadata were required as a selection criterion (date of isolate and country of origin was required) were also available. All public read sets were processed using the same local quality control thresholds mentioned previously and incorporated into the lineage specific analyses.

## Results

This study analysed 1,305 isolates spanning two decades from a total of 2,258 culture confirmed *Shigella* spp. cases reported in the state of Victoria. Two thirds (892/1,305, 68%) of the total number of *S. sonnei* were isolated from male cases (Table 1). However, this masks the significant shift in proportion of male case numbers 2014/2015.

**Table 1:**
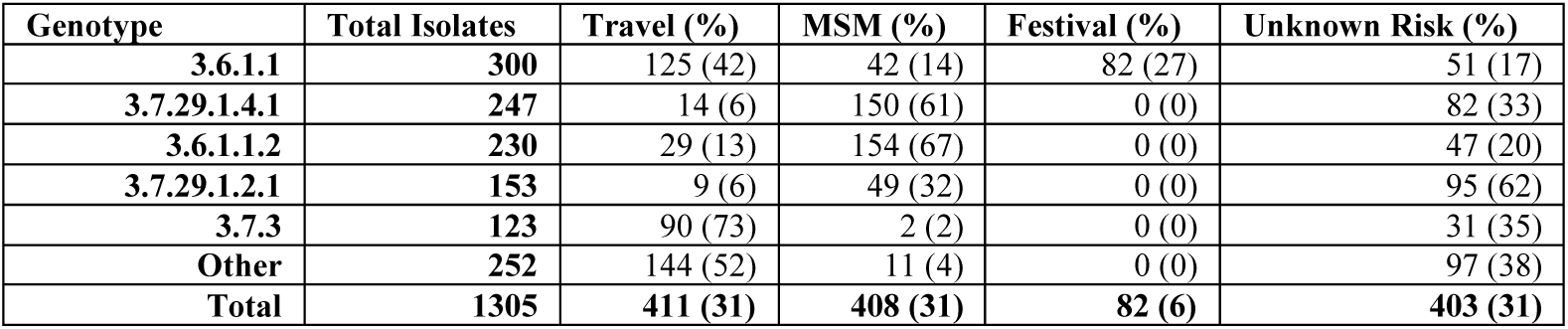
Dominant genotypes of *Shigella sonnei* isolates within Victoria and counts of isolates within each epidemiological risk group (Travel, Men who have sex with men (MSM), a large festival outbreak, and Unknown (where case not associated with a reported risk group or no risk was reported)

Three key epidemiological risk factors were identified, namely international travel (411/1,305, 31%), MSM-associated (408/1,305, 31%), and association with a large point source outbreak at an outdoor festival in early 2024 (82/1,305, 7%) (Table 1, Figure 1). The decrease in cases between 2020 to 2022 corresponds to SARS-COV-2pandemic and significant population-level interventions in place in Victoria, including border closures and lockdowns (Figure 1). In 2019, the main epidemiological risk factor changed from international travel to MSM-associated, and despite the drop in cases between 2020-2022, MSM-associated infections was the primary risk factor until the end of the study period in 2024. A total of 34 genotypes were detected in the dataset all of which were sub-lineages of global lineage 3^30^. Of note, five genotypes accounted for 80% (1,053/1,305) of the total dataset (Table 1, Supplementary Table 1). These five genotypes were found to be associated with a particular risk group (MSM or Travel) and were observed to fluctuate in prevalence over the surveillance period, with one genotype dominating in the Victorian the population at any one time. The broad AMR profiles at either MDR or XDR level changed with time. MDR was not detected prior to 2016 and remained in a low proportion of cases (6-11% cases per year) until 2020 where they comprised of 53% of cases (Figure 1). XDR was first detected in isolates from 2018; however, it was in 2024 when XDR cases increased to 69% of cases that year.

**Figure 1:**
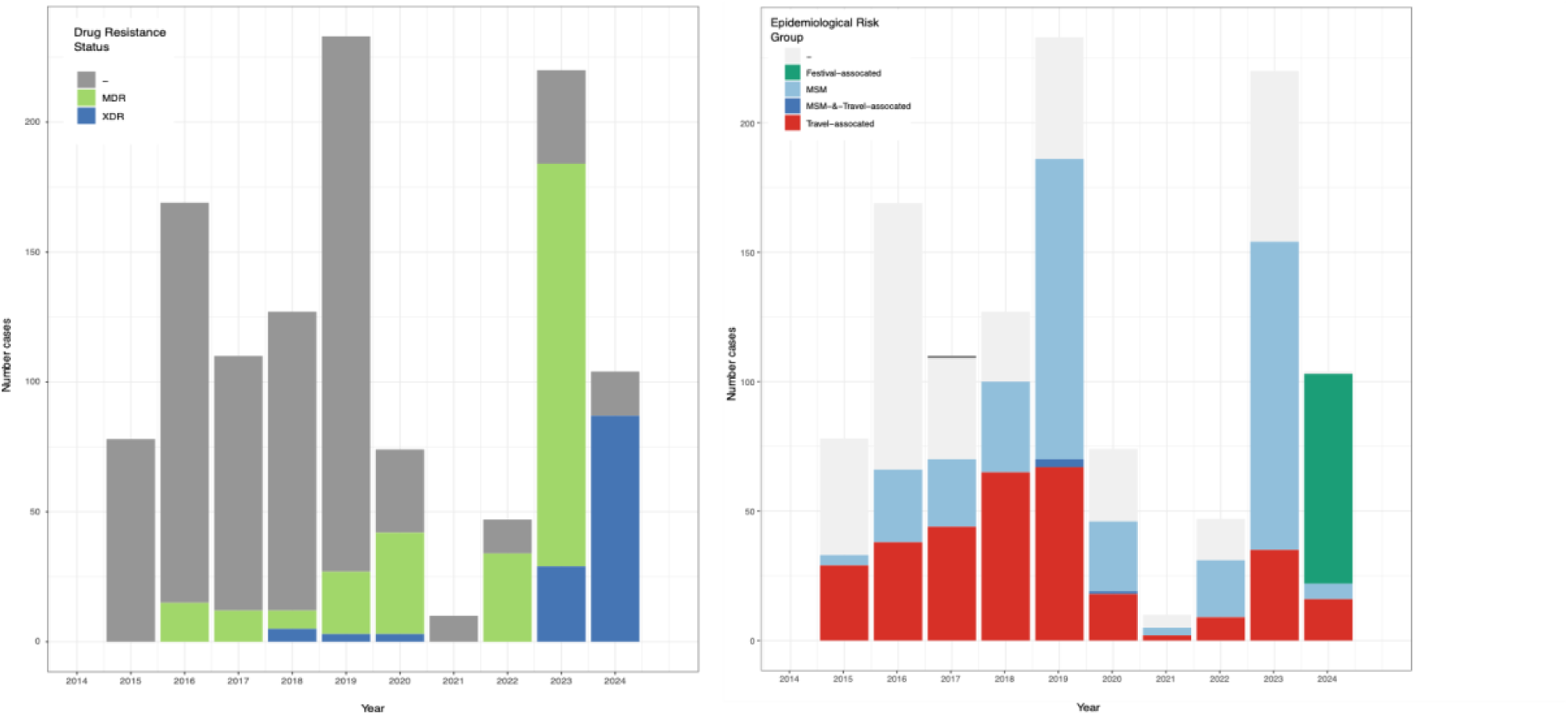
A. showing number of cases between 2015 and 2024 that were determined to be multidrug-resistant (MDR), extensively drug-resistant (XDR) or neither. B. showing number of cases between 2015 and 2024 with epidemiological risk factors; travel-associated, men who have sex with men (MSM), both travel and MSM, associated with an outbreak at a music festival, or no identified risk factor/not recorded.

Three independent Victorian sub-lineages of interest were observed; two sub-lineages were associated with MSM designated VIC-I, and VIC-II, and a third sub-lineage VIC-III associated with a festival outbreak (Figure 2), each associated with antimicrobial drug resistance. Genotyping in combination with phylogenetic placement confirmed that Victorian *S. sonnei* belonged to two lineages (3.6 and 3.7) (**Figure 2**). The large MDR lineage VIC-I was comprised of genotype 3.7.29.1.4.1 isolates that carried *bla*CTX-27 among other AMR genes to streptomycin, co-trimoxazole and macrolides (Supplementary information). The primary epidemiological risk factor for this group was MSM with limited international travel reported (Supplementary Table 1). The remaining two were defined as co-circulating XDR lineages, both genotyped as 3.6.1.1. However, the AMR profiles diverged between the two sub-lineages as did the primary epidemiological risk factor. One of these, VIC-III was linked to a large festival outbreak in 2024, with VIC-II linked to MSM as the primary risk factor.

**Figure 2:**
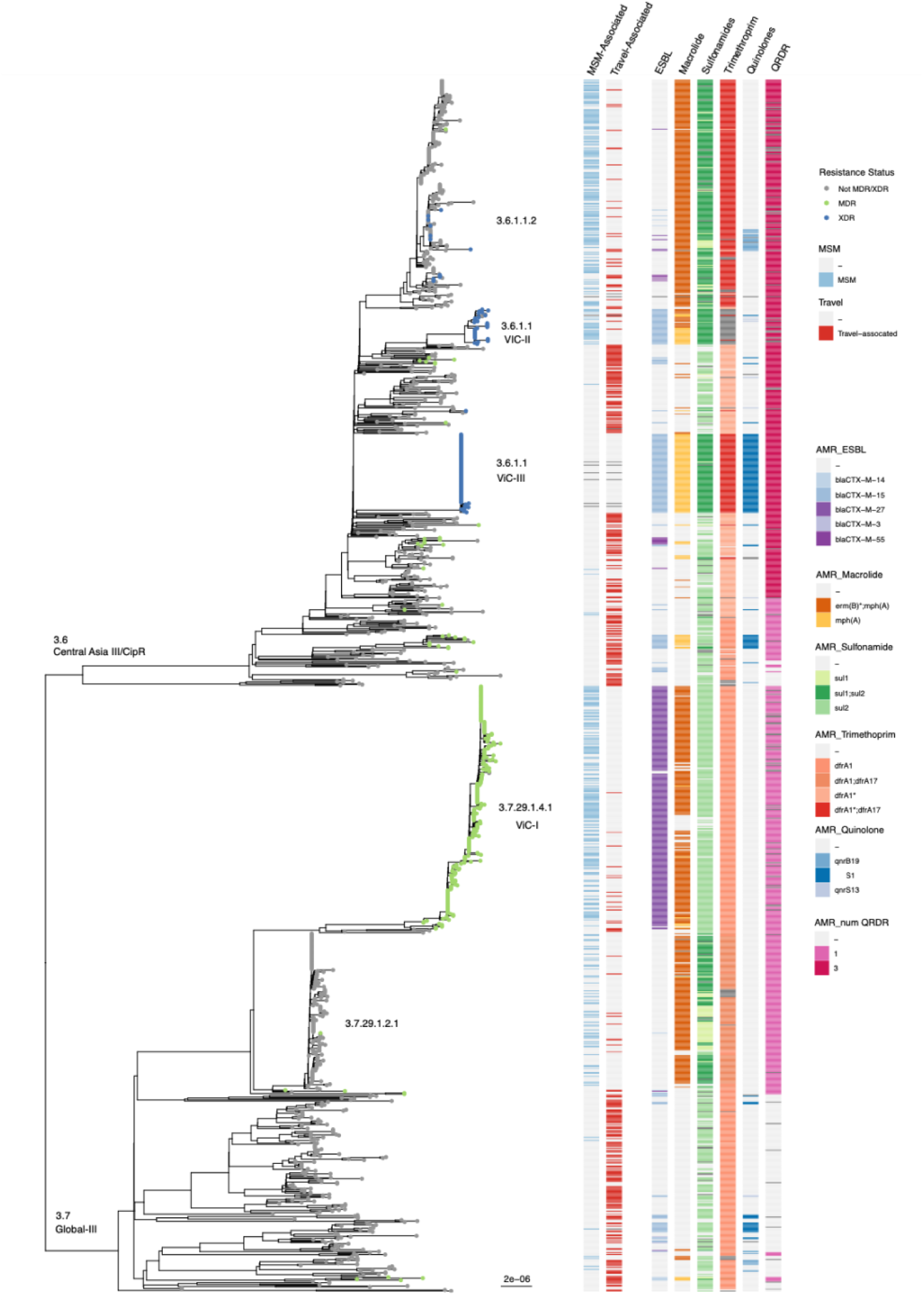
Maximum-likelihood phylogeny of 1,305 *Shigella sonnei* isolates from public health surveillance in Victoria, Australia from 2002-2024. The phylogeny was built from 6628 SNPs and 2790 parsimony informative sites using AUSMDU00008333 as the reference genome for mapping and SNP calling. IQtree2 was used to construct the phylogeny using the GTR+F+G4 model and 1000 rapid bootstraps. Tips of tree are coloured to reflect resistance status^16^ (grey neither MDR or XDR according to CDC definitions (REF), green multidrug resistance strains (MDR n=165) and blue extensively drug-resistant strains (XDR, n = 165). Heatmaps show the presence of primary risk categories and antimicrobial resistance determinants, where colours denote presence in that category and light grey being absent. Key lineages are denoted according to genotypes defined by sonneityper^25^ and lineages of interest VIC-I, II and III are labelled.

Travel-associated isolates were diverse (29 different genotypes) with multiple importation events into Australia. (**Supplementary Table 1**) The most common travel associated genotypes were 3.6.1.1 (42%, 125/411) and 3.7.3 (21%, 90/411), while the remaining 27 genotypes were less prevalent (1–29 isolates each). Among the 411 confirmed travel-associated isolates, 119 were non-specific in international travel, where location was unknown or multiple locations were visited, 9 involved travel only within Australia, and the rest were linked to 48 countries in 11 geographical regions (**Figure 3**). The most common regions were Southeast Asia (110 isolates) and South Asia (94 isolates). Isolates of lineage 3.7.3 were primarily from Indonesia (62 isolates), including 9 MDR and 3 XDR cases. Lineage 3.6.1.1 was most frequently associated with India (66 cases) and have mixed AMR profiles with 12 MDR and 2 XDR cases. MDR cases were mainly linked to travel in South and Southeast Asia, while XDR cases were found across multiple geographical regions. Phylogenetic and genetic distance analyses between isolates indicated that most travel-related cases were individual infections that were rarely associated with onward transmission. However, their introduction into high-risk transmission settings, such as MSM networks or large point-source outbreaks, could enable rapid local expansion and future outbreak potential.

**Figure 3:**
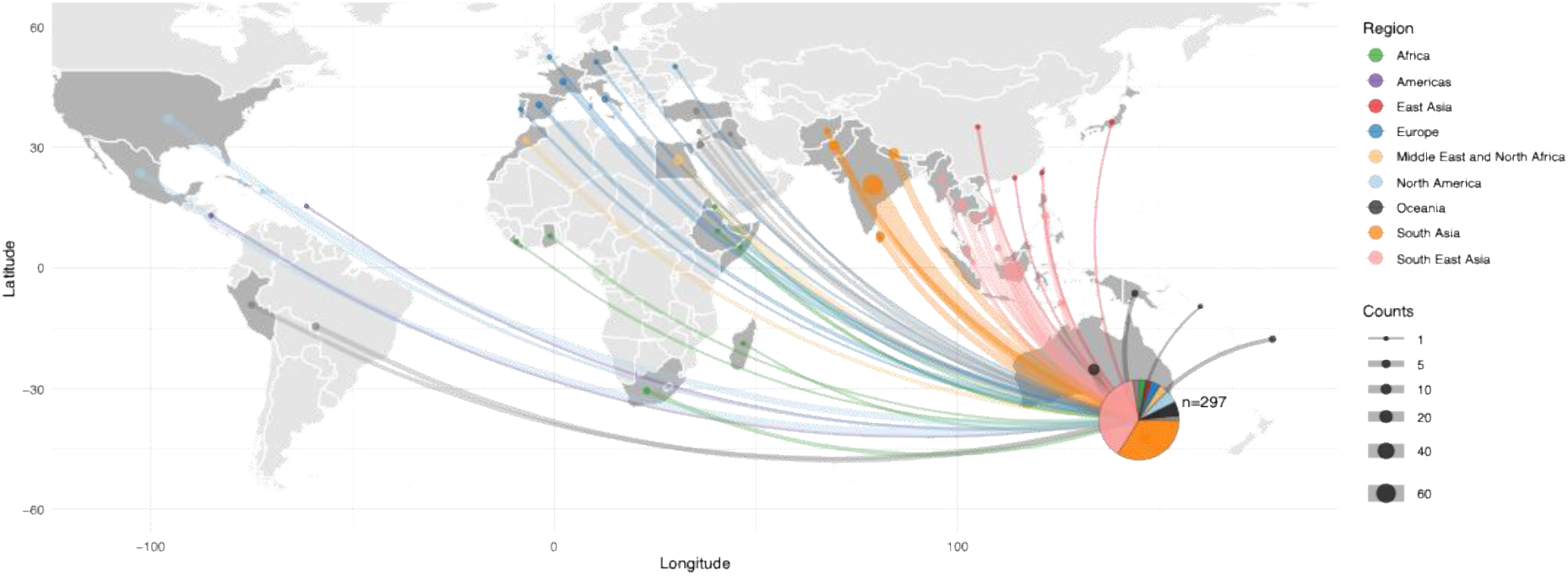
Distribution of *Shigella sonnei* travel-associated cases with regional breakdown. Map showing country/region of travel for *Shigella sonnei* travel-associated cases in Victoria, Australia. Colours refer to geographical region where case travelled, and countries where cases travelled to are shaded dark grey. Size of dot in each country is proportional to number of cases reporting travel to that country. Pie graph shows the breakdown of cases for travel to each geographical area.

In contrast to travel-associated cases, MSM-associated cases were genomically more homogeneous, with only seven genotypes, of which 44% were MDR (152/408) and 7.1% were XDR (29/408). Over the past decade, the dominant genotypes in Australia have changed (**Figure 4**), particularly in MSM-associated cases. Within this group, three genotypes followed a pattern of rapid expansion, dominance, and eventual decline, being replaced by new genotypes over time. For example genotype 3.7.29.1.2.1 (Global III VN2.MSM2.Aus) dominated MSM-associated cases from 2016 to 2017 (n=49/54, 91%). This genotype was subsequently replaced by 3.6.1.1.2 (MSM5) in 2018-2019 (n=137/154, 89%), before genotype 3.7.29.1.4.1 emerged as the dominant genotype from 2020 to 2023 (n=150/173, 87%). Of note, the expansion of this genotype was likely driven by the presence of *bla*CTX-M-27 resulting in resistance to 3GCs (**Figure 4**). Since 2024, a recently emerged subgroup of genotype 3.6.1.1 has been the most prevalent genotype circulating in the MSM population and is characterised by an XDR profile (**Table 2**). This continual genotype replacement corresponds to the persisting expansion of MDR and XDR profiles in these genotypes.

**Figure 4:**
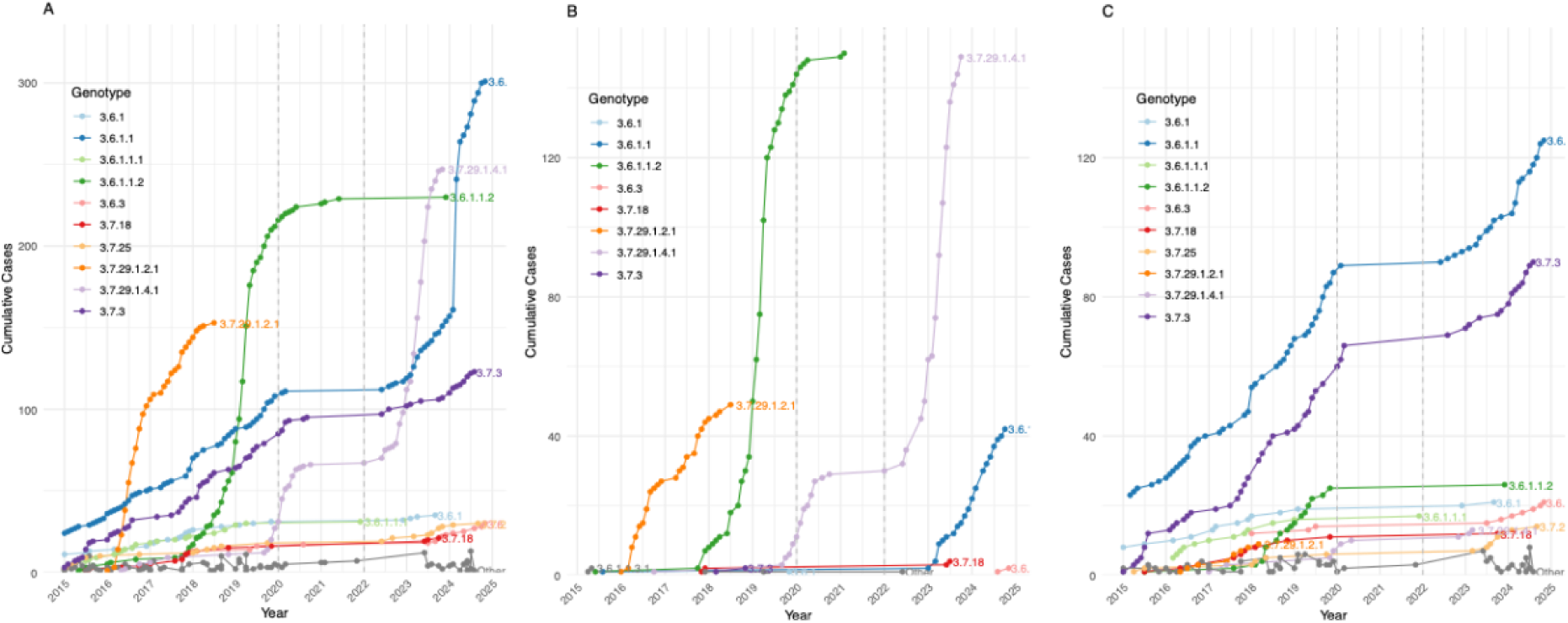
A. Timeline accumulation plot of the *S. sonnei* genotypes of B. MSM-associated genotypes and C. international travel-associated genotypes between 2015 and 2024. Top 10 most common genotypes are coloured.

**Table 2:** Antimicrobial profiles of Victorian sub-lineages of interest: Full matches (100% protein match to AMR gene in database) and close matches (>90% of a gene was recovered with >90% protein identity to a gene in the database. Close matches are reported in the output files with an asterisk (e.g. ermD*) at those with a ^ indicate a partial gene match.

|  | Clinically relevant antimicrobial resistance |  |  |  |  |  | Other antimicrobial resistance |  |  |  |
| --- | --- | --- | --- | --- | --- | --- | --- | --- | --- | --- |
|  | ESBL | Macrolide | Sulfonamide | Trimethoprim | Quinolone | QRDR | Streptomycin | Efflux | Tetracycline | other |
| VIC-I | *blaCTX-M-27* | *erm(B)\*;mph(A)* | *sul2* | *dfrA1* | - | *gyrA_S83L* | *aadA1;aph(3'')-Ib;aph(6)-Id* | *emrD\** | *tet(A)* | - |
| VIC-II | *blaCTX-M-15* | *erm(B), mph(A)* | *sul1;sul2* | *dfrA1\*;dfrA17* | *qnrS1* | *parC_S80I, gyrA_S83L, gyrA_D87G* | *aadA1;aadA5* | *emrD\*;acrF\*^* | *tet(A)* | *sat2* |
| VIC-III | *blaCTX-M-15* | *mph(A)* | *sul1;sul2* | *dfrA1\*;dfrA17* | *qnrS1* | *parC_S80I, gyrA_S83L, gyrA_D87G* | *aadA5;aph(3'')-Ib\*;aph(6)-Id* | *emrD\*;acrF\*^* | *tet(A)* | *sat2* |

Three recent sub-lineages of concern were identified in these data, characterised as significant public health threats due both to their size and/or drug-resistance profile. The largest sub-lineage VIC-I contained 247 isolates of genotype 3.7.29.1.4.1 (Global III VN2.KH1.Aus) carrying *bla*CTX-M-27 and classed as MDR. 221 genomes clustered <5 SNPs apart (within species phylogeny), spanned the five years (2017–2023), only to then disappear in 2024. Among VIC-I, 92% (229/247) of isolates came from men that were aged between 18-50. An epidemiological risk of MSM-association could be confirmed in 61% (150/247). This prolonged outbreak was characterised by 3GC resistance; 98% (243/247) isolates within this sub-lineage carried ESBL gene *bla*CTX-M-27, and were classified as MDR with resistance mechanisms detected for macrolides, co-trimoxazole and aminoglycosides^12^. Isolates were ciprofloxacin susceptible with *gyrA* (S83L) being the only mutation observed within the QRDR.

VIC-I increased in prevalence in 2019 to become the more dominant sub-lineages in the Victorian MSM risk group (**Figure 5**), reported cases stalled throughout the SARS-CoV-2 pandemic. After the removal of public health measures, VIC-I rapidly expanded locally within the MSM population. We quantified the SNP accumulations over time within the VIC-I sub-lineage. A range of 0-60 pairwise SNPs was identified between VIC-I strains, with an estimated accumulation of 0-15 SNPs per year across the genome. Comparing isolates before and after the pandemic, 11 SNPs differed between the two periods. The SNP accumulation in combination with no international isolates intermixing between this period suggest that VIC-I may have persisted in Victoria throughout the SARS-CoV-2 pandemic restrictions, accumulating SNPs within the expected range (0-15 per year). (**Figure 5**). Incorporation of 187 global context isolates (genome detection portal cluster PDS000019750.343) indicated a potential common ancestor of VIC-I to be from MSM cases in Canada 2017^21^. After the post-pandemic expansion of VIC-I in Victoria, there were indications that this lineage was subsequently exported from Australia as several closest relatives of recent international cases in the USA are this same sub-lineage.

**Figure 5:**
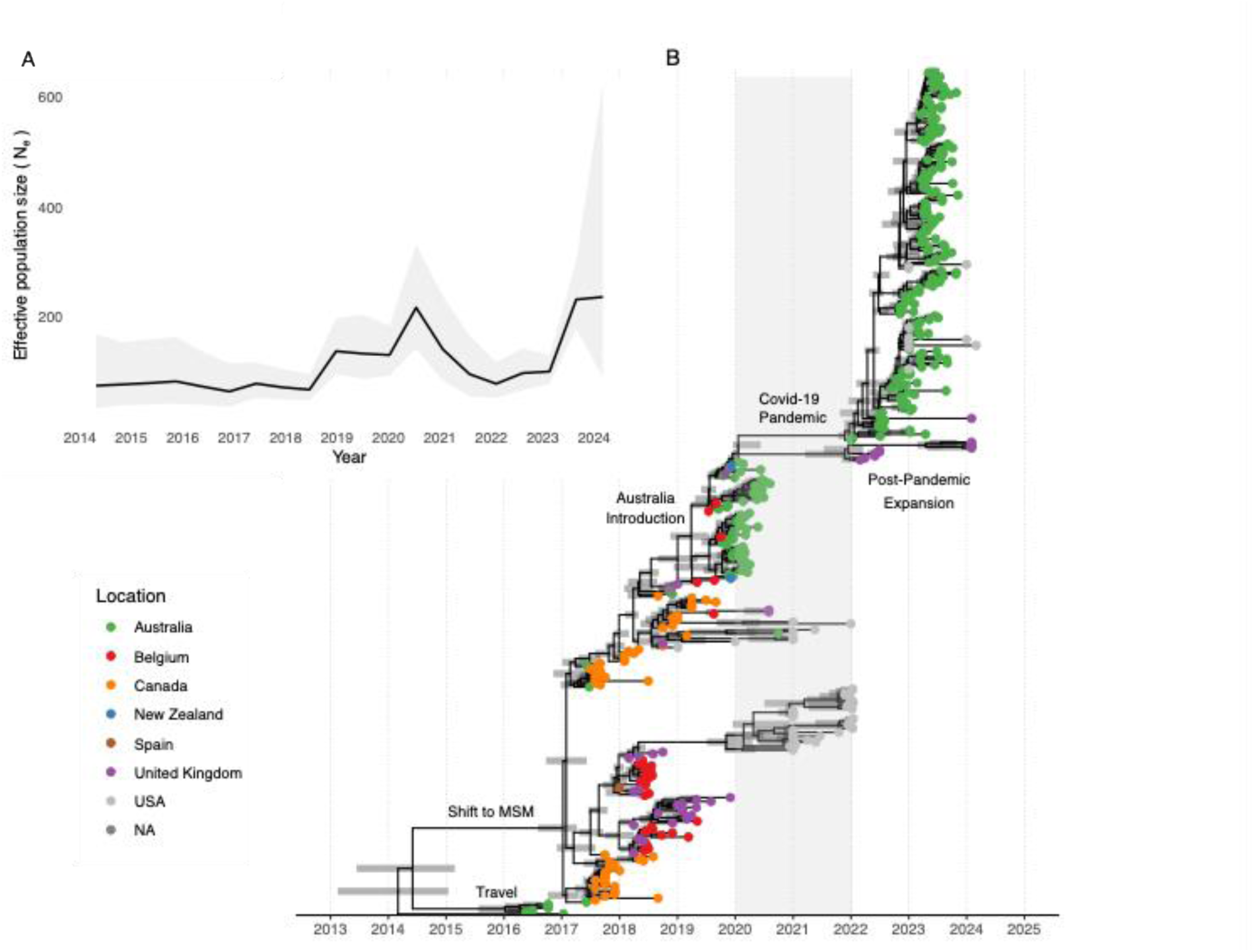
Maximum-likelihood time-based phylogeny of 245 *Shigella sonnei* 3.7.29.1.4.1/MDR VIC-I sub-lineage in Victoria, Australia from 2016-2024 with global context isolates. A) Skyline plot showing changes in effective population size Ne over time, inferred through ancestral state reconstruction of genetic data. B) The phylogeny was constructed using 245 SNPs, including 103 parsimony-informative sites, with AUSMDU00022573 used as the reference genome for read mapping and SNP calling. The tree was inferred using IQ-TREE2 with the GTR+F+G4 substitution model and 1,000 rapid bootstrap replicates. TreeTime was used to calibrate the phylogeny in time by converting internal nodes and tips to estimated dates. Tips are coloured according to the country of isolation. Horizontal bars represent the 95% confidence intervals for node date estimates. Key inferred events are indicated along the timeline, including the shift to MSM-associated transmission, introduction into Australia, and the period of pandemic impact.

MSM sub-lineage VIC-II of *S. sonnei* was characterised as XDR through phenotypic testing. VIC-II was defined as isolates within a monophyletic subgroup of the genotype 3.6.1.1 (aka CipR) within Australian context (**Figure 2**), however in the global context represents a polyphyletic group (**Figure 6**). Two long branches in the phylogeny separated both sub-lineages from all other 3.6.1.1 isolates in Victoria (**Figure 6**). All VIC-II isolates carried the ESBL gene b*la*CTX-M-15, plus macrolide, trimethoprim, and aminoglycoside resistance genes and the triple *gyrA* (S83L), *gyrA* (D87G) and *parC* (S80I) mutations in the QRDR leading to high-level resistance to ciprofloxacin. VIC-II contained 53 isolates and were all detected in Victoria in 2023-2024. 82% came from men, amongst which MSM association could be confirmed in 66% (35 isolates), while two had recent travel to USA and Thailand. Within the Victorian dataset VIC-II formed a unique sub-lineage of genotype 3.6.1.1 isolates, with a long branch length and far genetic distance with median 70 (IQR 55-80) SNPs to all other 3.6.1.1 isolates forming a tight monophyletic clade. Global contextualisation with 190 publicly-available genomes revealed potential origins of this clade from the United Kingdom^31^ (earliest sequence from April 2022), which also suggests that the VIC-II sub-lineage has been likely introduced into Victoria on multiple occasions after border restrictions eased, forming a polyphyletic group with the United Kingdom isolates.

**Figure 6:**
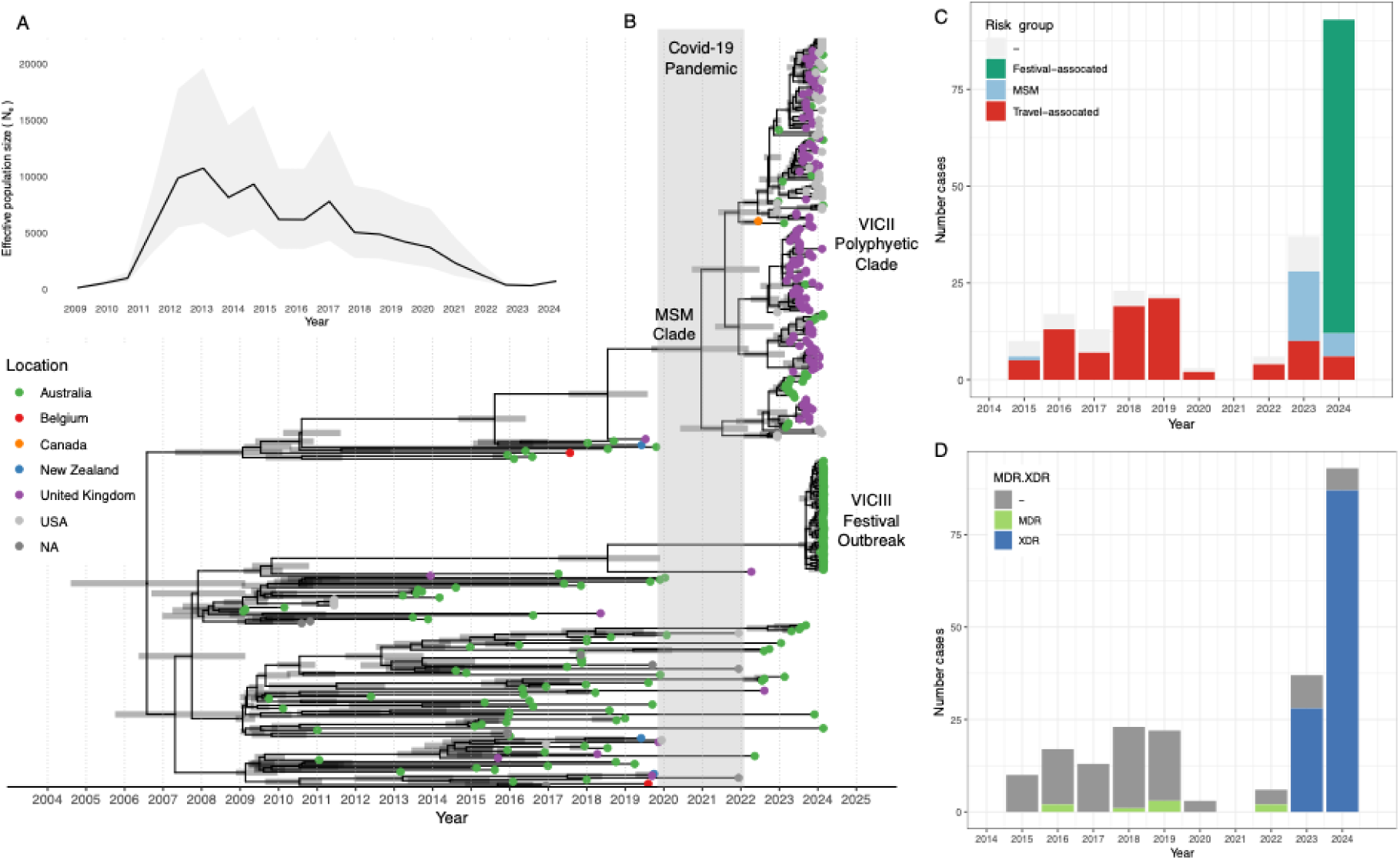
Maximum-likelihood time-based phylogeny of 248 *Shigella sonnei* 3.6.1.1 sequences, including the XDR VIC-II and VIC-III clades in Victoria Australia from 2016-2024 withing global context samples. A) Skyline plot showing changes in effective population size (Ne) over time, inferred through ancestral state reconstruction of genetic data. B) The phylogeny showing clear separation of VIC-II and VIC-III from other 3.6.1.1 isolates sub-lineages was built from 1887 SNPs and 516 parsimony informative sites using AUSMDU00010908 as the reference genome for mapping and SNP calling. IQtree2 was used to construct the phylogeny using the GTR+F+G4 model and 1000 rapid bootstraps. TreeTime was used to transform tips and nodes to dates. Tips are coloured according to the country of isolation. Horizontal bars represent the 95% confidence intervals for node date estimates. Key inferred events are indicated along the timeline, including the shift to MSM-associated transmission, introduction into Australia, and the period of pandemic impact. C. Number of cases between 2015 and 2024 that were determined to be multidrug resistant (MDR), extensively drug resistant (XDR) or neither. D. Number of cases between 2015 and 2024 that were attributed to travel, MSM risk or neither group.

Isolates from the festival outbreak sub-lineage VIC-III were classified as XDR, with an AMR profile that differs only slightly from VIC-II due to the absence of the *ermB* gene, and varied streptomycin resistance genes (*aph(3’’)-Ib*;aph(6)-Id*, instead of *aadA1*) (**Table 2**). VIC-III represents a distinct branch within the genotype 3.6.1.1 and biotype f, distinct from the VIC-II sub-lineage (**Figure 6**). Eighty-two genomes were attributed to 65 individual cases epidemiologically associated with a lifestyle festival outbreak (0-3 pairwise SNPs between cases). Comparing the outbreak isolates with global genomes, no closely related isolates were identified from Victoria or publicly available sequences. This suggests that VIC-III may have been introduced from an unknown location and followed by rapid spread through the festival. Transmission at this event is not considered to be connected to MSM behaviour (62% female). It is suspected that an initial point-source transmission event occured, which then led to rapid transmission amongst staff and patrons exacerbated by high temperatures, an outdoor camping environment, potentially poor hygiene measures (including limited shower facilities and not washing hands thoroughly), shared toilet facilities and extensive close contact without the ability to isolate when unwell.

## Discussion

In this study we described the population dynamics of *Shigella sonnei* in Victoria Australia over two decades. Within the men who have sex with men (MSM) population, a pattern of rapid expansion and clonal replacement was observed over the past decade. Throughout the surveillance period we observed four lineage replacements within MSM communities, whereby each sub-lineage was found to persist for approximately two years (excluding pandemic impacts), before reducing in prevalence and being overtaken by a new lineage a trend being previously indicated in other countries^32^. This turnover coincides with the yearly fluctuations of cases within Victoria, and with an alarming trend toward increased AMR resulting in a dominant XDR genotype in circulation (Figure 4).

The MDR lineage VIC-I has been a concern since its first description^13^ with cases rising through 2019-2020. During the SARS-CoV-2 pandemic^33^, restrictions in Victoria reduced social movement and slowed expansion but cases surged again post-restrictions suggesting only limited disruption to *Shigella sonnei* transmission. Genomic data confirm persistence of this lineage throughout, rather than re-introduction with consistent SNP variation across timepoints. Post-pandemic VIC-1 case numbers peaked to levels comparable with the previously dominate strain before rapidly declining, with only one case detected in 2024. This pattern suggests that once ∼150 isolates of a genotype are observed through the state public health laboratory, transmission may reach a saturation point within the affected population. Such dynamics, seen in pathogens like E. coli ST131^34^, likely reflect constrained susceptible host pools, transient immunity, or strain ‘burn-out’, highlighting how transmission can self-limit even in highly connected communities.

A particularly concerning development however is the emergence and establishment of the XDR VIC-II lineage. Its rapid appearance towards the end of 2023 in the MSM community alongside the reduced cases of VIC-I is concerning. If VIC-II follows the trend of the previous MSM dominant sub-lineages, it poses significant potential harm due to its expanded resistance, reducing treatment options. Monitoring these concerning genotypes is crucial to managing health risks and preventing widespread transmission as this sub-lineage already appears to be commonly isolated in the United Kingdom^17^ and similar trends have been reported in France^19^. Additionally this trend has also be observed in *Shigella flexneri*^21^ for which cases are also increasing^35^.

In contrast to MSM-associated cases, most travel-related infections in Victoria were sporadic and did not lead to sustained outbreaks. However, the major MSM lineages VIC-I and VIC-II were likely introduced via international travel (e.g., Canada-linked VIC-I in 2017^36^ and UK-linked VIC-II in 2023^1^) and subsequently established within highly connected local networks. While most travel-associated cases are low risk, introductions into such populations can enable rapid local spread^13,37^. Victoria also appears to contribute to global dissemination, with genomic evidence suggesting export of VIC-I to the USA post-pandemic^38^. Together, these findings show that MDR and XDR Shigella sonnei transmission is not a local issue but part of an interconnected global network. This underscores the need for coordinated international surveillance and open genomic data sharing to accurately track strain movement and distinguish local transmission from global spread.

The high level of global exchange of MDR and XDR *Shigella* within MSM populations emphasised in our study and multiple others^9,10,21,31^, changes the perspective on public health surveillance of *Shigella*, from a localised jurisdictional issue to a shared global problem^20,39^. Because *S. sonnei* circulates globally, open data sharing is essential for resolving strain origins and distinguishing local evolution from international introductions Crucially, without publicly available datasets and the use of pathogen detection portal, the scale and pathways of global spread evident in our results would have possibly misattributed to local circulation instead of part of a global dissemination.

Towards the end of March 2024, a large outbreak of XDR *S. sonnei* occurred at a festival. Pre-existing genomic surveillance within Victoria of *S. sonnei* enabled the rapid identification of this new sub-lineage VIC-III as unlinked from local and international isolates. While distinct from all other Victorian isolates this outbreak raised further concerns about the antimicrobial status of *S. sonnei*. There are now two clades of XDR within Victoria of the 3.6.1.1 which have impacted distinct populations. Horizontal gene transfer between strains has been shown to be the cause of new emergent genotypes^14,20,40^ with increased AMR so the presence of multiple XDR genotypes in the Victorian population increases that risk, genotypes previously considered non-threatening now have a greater chance of acquiring new resistance genes, due to the larger genetic pool of available resistance genes.

This study has several limitations. We did not assess *Shigella sonnei* dynamics alongside other co-circulating enteric or sexually transmitted pathogens. Culture-confirmed cases underrepresent PCR-notifications^6^, so population dynamics may be incomplete despite enhanced surveillance. Metadata gaps particularly pre-2015 MSM risk data and limited travel information restrict source attribution. The analysis is confined to a single state, while transmission often spans jurisdictions leading to multi-state outbreaks. We also did not assess asymptomatic carriage or close contacts, limiting links between genotypes, disease severity, and transmission risk.

*Shigella sonnei* sits at the intersection of foodborne disease, sexually transmitted infections, and antimicrobial resistance, yet surveillance systems often treat these domains separately, risking missed transmission links and delayed detection of emerging MDR/XDR lineages. Our findings in this study demonstrate the value of genomic epidemiology for high-resolution genotyping, AMR profiling, and tracking of high-risk introductions, while highlighting the need for integrated surveillance across enteric, sexual health, and AMR frameworks. Strengthening coordinated international data sharing and systematic reporting will be essential to accurately track transmission and respond effectively to this increasingly complex global public health threat.

## Acknowledgments

MDU PHL is funded by the Department of Health, State Government of Victoria. We would like to acknowledge the diagnostic laboratories across Victoria for submitting isolates. MDU PHL laboratory staff for their role in sample testing, analysis and reporting. We thank the staff of the Department of Health Victoria and Local Public Health Units for their support, providing epidemiologic data used in this analysis, and assistance with interpretation.

