## Supplementary Information for "Incremental AMR acquisition driving successive genotype replacements and the rise of extensively drug resistant (XDR) *Shigella sonnei* in Australia over 20 years"

### Supplementary Figures

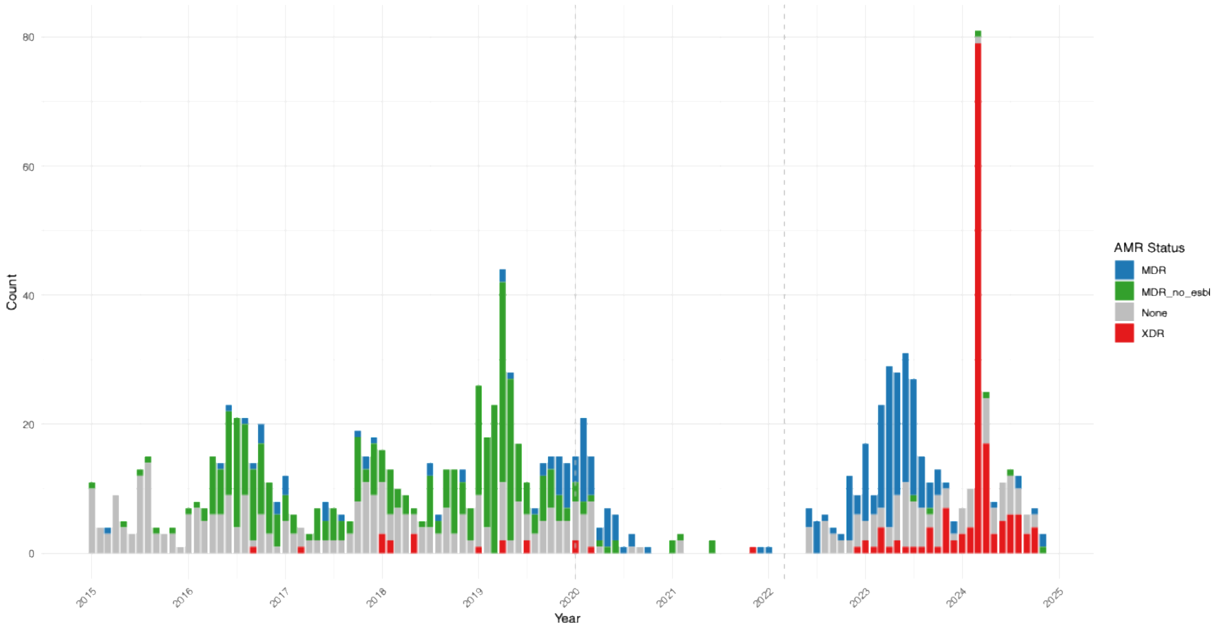


**Supplementary Figure 1:** Epidemiological plot showing monthly case counts of Victorian *Shigella sonnei (post-2015)* coloured by AMR status as determined by the CDC definitions. Multidrug resistant strains (MDR) = green, MDR with extended beta lactamase genes (ESBL) = blue and extensive drug resistance (XDR) = red, grey = neither MDR or XDR strains.

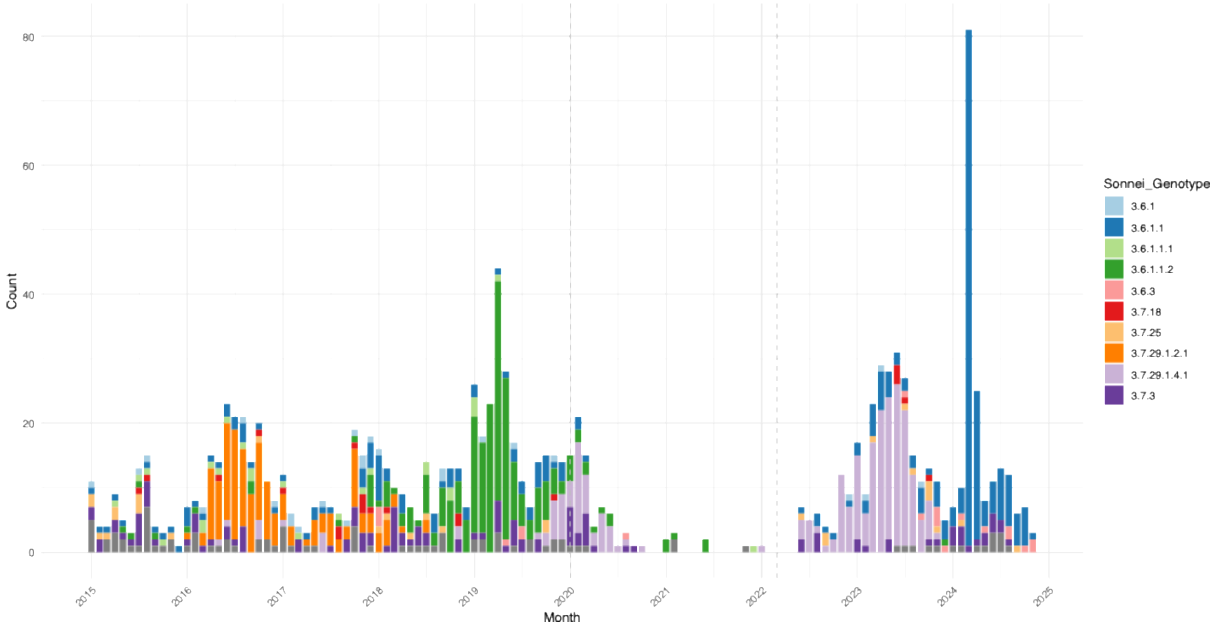


**Supplementary Figure 2:** Epidemiological plot showing monthly case counts of Victorian *Shigella sonnei (post-2015)* coloured by the 10 most prevalent lineages as determined with sonneityper.

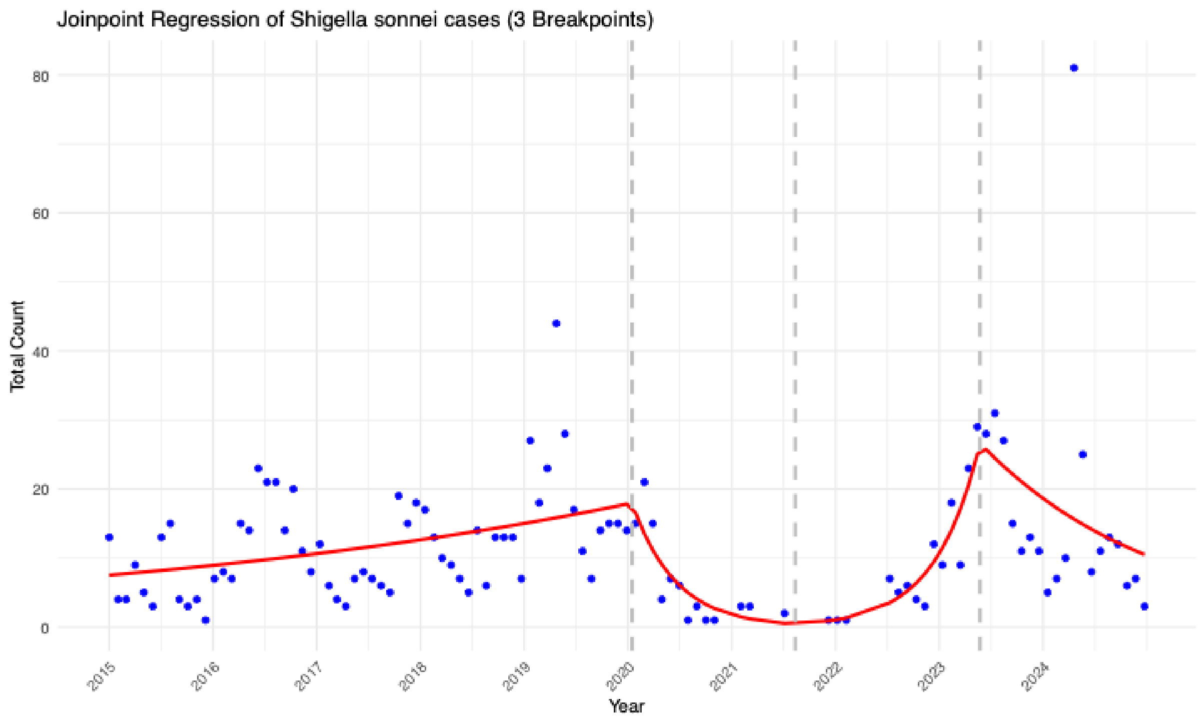


**Supplementary Figure 3:** Joinpoint regression showing monthly case counts of Victorian *Shigella sonnei (post 2015).* Three breakpoints shows a shift in case trends around the times of covid interventions in Victoria.

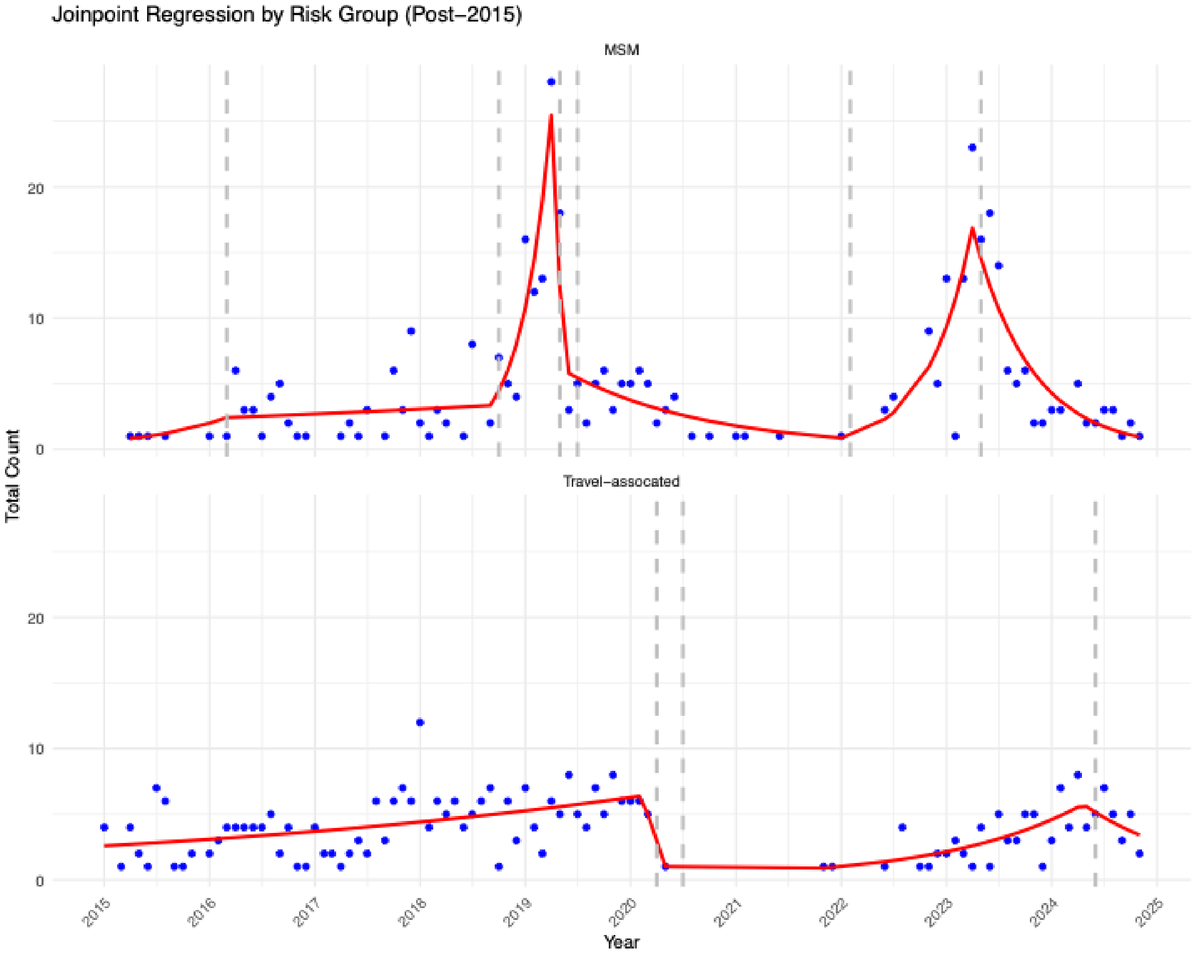


**Supplementary Figure 4:** Joinpoint regression showing monthly case counts of Victorian *Shigella sonnei (post 2015) faceted* by risk group. Six breakpoints were associated with MSM cases while only three breakpoints were associated with Travel.

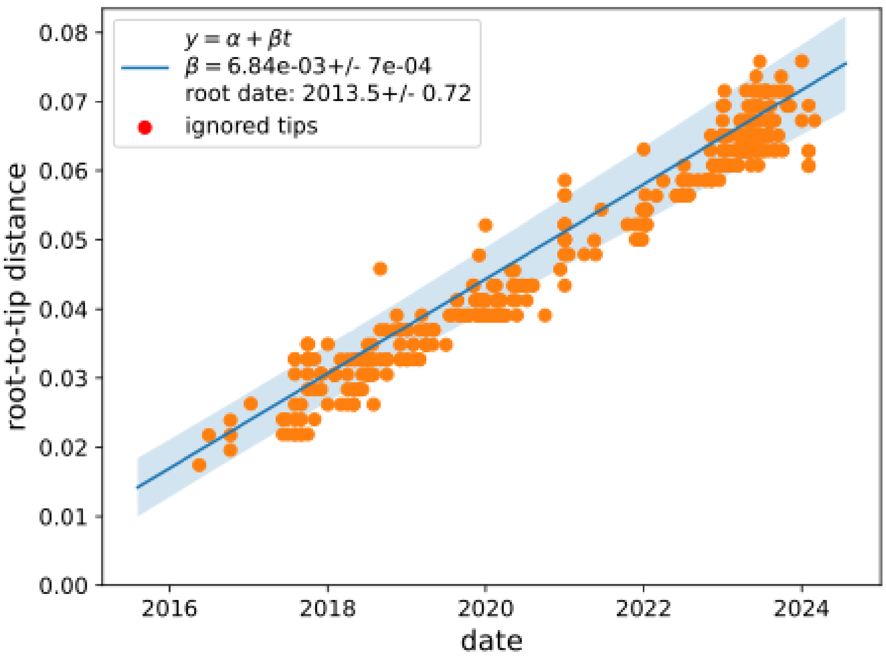


**Supplementary Figure 5 :** Root to tip regression of maximum-likelihood tree for lineage 3.6.1.1

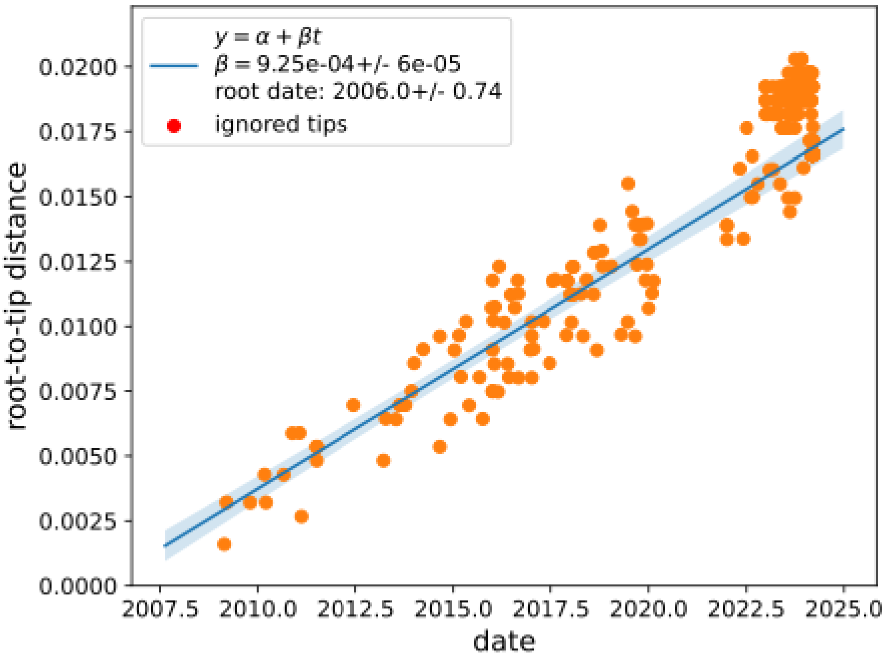


**Supplementary Figure 6 :** Root to tip regression of maximum-likelihood tree for lineage 3.7.29.1.4.1
